# Loss of Photosynthetic RhythmThermal Plasticity Under Domestication and Repurposing Drivers of Circadian Clock (DOC) Loci for Adaptive Breeding in Barley

**DOI:** 10.1101/2020.05.15.098418

**Authors:** Manas R. Prusty, Eyal Bdolach, Eiji Yamamoto, Jeffrey L. Neyhart, Lalit D. Tiwari, Klaus Pillen, Adi Doron-Feigenbaum, Kevin P. Smith, Eyal Fridman

**Affiliations:** Institute of Plant Sciences, Agricultural Research Organization (ARO), The Volcani Center, P. O. Box 6, 5025001, Bet Dagan, Israel; Department of Life Sciences Faculty of Agriculture, Meiji University, Japan; Department of Agronomy and Plant Genetics, University of Minnesota, St. Paul, MN 55108; Institute of Agricultural and Nutritional Sciences, Martin-Luther University Halle-Wittenberg, Halle (Saale), Germany

**Keywords:** Circadian clock, Domestication, Plasticity, Barley, Genome-wide association studies, Crop yield, QTL-environment association, Local adaptation

## Abstract

Circadian clock rhythms are critical to control physiological and development traits, allowing, plants to adapt to changing environments. Here we show that the circadian rhythms of cultivated barley (*Hordeum vulgare*) have slowed and amplitude increased under domestication by comparing with its wild ancestor (*H. spontaneum*). Moreover, we show a significant loss of thermal plasticity during barley evolution for the period and more extensively for amplitude. Our genetic analysis indicates that wild allele at epistatic loci, which mutually condition clock variation and its thermal plasticity in interspecific crosses, are absent in a contemporary barley breeding panel. These epistatic interactions include conditioned effects of Drivers of Circadian (DOC) clock loci on chromosome 3 and 5, which mediate amplitude decrease and period lengthening, respectively, under domestication. Notably, two significant loci, *DOC3.1* and *DOC5.1*, which are not associated with clock diversity in cultivated breeding material, do show pleiotropic effects on flowering time and grain yield at multiple experimental sites across the U.S. in a temperature-dependent manner. We suggest that transition from winter growth of wild barley (*H. spontaneum*) to spring growth of modern cultivars included the loss and repurposing of circadian clock regulators to yield adaptation by mechanisms yet to be clarified.

**Significance statement:** Circadian clock rhythms are crucial factors affecting crop adaptation to changing environments. If faced with increased temperature plants could respond with temperature compensation adaptation and maintain clock rhythms, or they can change period and/or amplitude to adapt. We used a combination of approaches: high-throughput clock analysis under optimal and elevated heat conditions, genome-wide association study (GWAS) with cultivated and wild diversity panels to identify changes under domestication and quantitative trait loci (QTL) that control the clock and its responses, and QTL-environment association for testing environmentally-conditioned effects of these QTLongrain yield and flowering timingacross US. Our findings provide insights into changes of circadian rhythms under domestication and genetic tools for plant breeders to develop better-adapted cultivars to changing environments.

## Introduction

Growth and metabolism are following rhythms that allow a resonance between environment dynamics, e.g. day and night, and molecular pathways that regulate these biological activities. The core circadian clock is what drives these rhythms, and it is reflecting on the cyclic pattern of different layers or outputs such as photosynthesis, cell division, and metabolism, e.g. starch synthesis and degradation. One hallmark attribute of the circadian clock machinery is that it maintains a relatively stable cycle of about (circa) 24 hours (dies)(1). Nevertheless, one emerging question in the study of the circadian clock is to what extent is it robust to environmental changes and does plasticity rather than such robustness or temperature compensation has been selected during natural or artificial evolution, i.e. crop domestication. The change in characteristics of the circadian clock has a major influence on the growth of plants and synchronization with the diurnal cycle is considered a critical adaptive feature of the clock (2, 3). Nevertheless, few studies have utilized naturally occurring variation (4), or made a systematic comparison between wild and cultivated plant material to realize if buffering or flexibility of the circadian clock against increased temperature is more beneficial at the population level. Alternatively, it is not clear yet if selection under domestication and breeding worked against or for plasticity of the clock, and if so, to what characteristic of the clock (period or amplitude, or both)?

The core clock in plants composed of three interlocked feedback loops, in which two Myb domain transcription, CIRCADIAN AND CLOCK ASSOCIATED1 (CCA1) and LATE ELONGATED HYPOCOTYL (LHY), are the hubs in these loops. The day-time expression of CCA1 acts to suppress expression in two loops, of TOC1 gene and of the evening complex including ELF4, LUX, GI and ELF3, which in turn work to suppress CCA1 and LHY during the night. In parallel, LHY and CCA1 protein complex are promoting the expression of a family of pseudo-response regulator (PRR) gene family, which in turn close the third loop and suppress LHY and CCA1 to facilitate the day-night cyclic expression of the core clock genes (1). Besides these three interlocked feedbacks loops, the core clock includes additional genes that affect the clock by posttranslational modifications and stabilization of its core components, dependent and independent of the light quality and quantity. The F-box protein ZEITLUPE (ZTL) and clock element GIGANTEA (GI) heterodimerize in the cytosol and hold the later from entering the nucleus and promote flowering (5). Notably, such modifications are rhythmic and also act on gene products in physiological, metabolic and signaling pathways (6), which consider as outputs of the core circadian clock machinery. Photosynthetic activity is one such output that became relevant for the development of non-invasive high throughput measurement of the circadian clock. These remote methods are including prompt and delayed fluorescence (F and DF) from Chl(7–9).

Transcript and protein abundance of circadian rhythms have been observed in barley plants as well (10–13). Several studies have reported the diurnal and circadian expression for HvLHY (HvCCA1), HvPPD-H1, HvPRR73, HvPRR59, HvPRR95, HvGI, HvTOC1, HvLUX and HvELF3(14–19). By mutant analysis, three barley clock genes have been well characterized: Ppd-H1, ELF3 and LUX (15, 18–20). The two early maturity mutants, early maturity 8 (*eam8*) and *eam10* barley genes were homologs of the Arabidopsis clock genes ELF3 and LUX1, respectively and mutants plants of both genes showed photoperiod insensitivity and early flowering under long- and short-day conditions in barley (18–20). More recently, analysis of diel and circadian leaf transcriptomes in the cultivar Bowman and derived introgression lines harboring mutations in EARLY FLOWERING 3 (ELF3), LUX ARRHYTHMO 1 (LUX1), and EARLY MATURITY 7 (EAM7) allowed Muller et al. (21) to predict the structure of the barley circadian oscillator and interactions of its components with day-night cues. In fact, some of the critical phenotypes under crop domestication were linked with mutations in the circadian clock genetic network and connected between clock variation to life-history traits relevant for adaptation to agricultural set-up and changing relevant traits (see detailed review (22). In wheat, barley and sorghum PRR-mediated insensitivity to long days was crucial for the transition to the Northern hemisphere, yet it was not implicated in life cycle control but more in the regulation of photoperiod flowering (15, 23). At the same time, selection for other clock gene alleles, e.g. *eam8* mutated at the barley HvElf3, which disrupt the dependency of flowering by time-of-the-day light inputs, allowed cultivation further north from the Fertile Crescent, in the shorter growing season of Europe (19). Signature of selection under domestication for clock traits and genes underlying were also reported recently for tomato, in which mutations in *LNK2* and *EID1*, both involved in light input to the circadian clock, were favored in the cultivated varieties (24).

In this study, we utilized our developed SensyPAM high-throughput F-based circadian rhythm measurement platform to estimate and compare the rhythmicity of the clock. We performed this phenotypic analysis on three barley populations with varying proportions of wild vs cultivated allelic diversity. We performed a genome scan, including single and two-dimensional GWAS to describe Drivers of clock (DOC) loci underlying the changes in circadian clock characteristics (their plasticity) between optimal and high temperature environments. Our analysis indicates the significant changes in rhythmicity including period and amplitude, as well as loss of plasticity between the two environments, has occurred underdomestication. Furthermore, comparison between genomic architectures underlying clock variation in wild and cultivated populations point to the loss of alleles at specific loci for clock deceleration and for its thermal plasticity. Finally, a QTL–climate association analysis between DOC loci and field phenotype across US indicate the repurposing of plasticity clock QTL for determining grain yield and heading variation thus revealing their potential for adaptive breeding.

## Results

### Hordeum vulgare populations with reduced levels of wild H. spontaneum DNA diversity show clock deceleration, increased amplitude and reduced thermal plasticity

Our initial goal was to compare populations with varying proportions of wild vs cultivated alleles for their thermal circadian clock plasticity. To estimate the rhythms of the clock we used the high throughput F analysis of plants at the 3-4 leaf stage, after these were entrained under long days or short days (14 h of light and 10h of dark and vice versa; see Methods). We estimated the period and amplitude of the clock rhythms from non-photochemical quenching (NPQ) levels measured under continuous light for 3 days. These experiments were conducted under optimal temperature (OT; 22°C) and high temperature (HT; 32°C), which allowed us to calculate the period and amplitude thermal plasticity, i.e. the change of the mean value of the trait for each line between the two environments (HT-OT). Previously we used this SensyPAM platform to extract these clock parameters from a biparental wild barley doubled haploid population (8) however, since we wished to obtain a better representation of the wild gene pool we expanded our analysis. We included representative accessions that were collected in each of the micro-sites of the wild barley infrastructure, Barley1K (B1K), and encompassed all the 51 sites that represent a broad genetic and ecogeographic adaptation (25). On average, the period of the B1K accessions was shortened under HT (HT-OT) by -1.6 hr (Fig. 1a). Nevertheless, there was a varying level of responses to heat between accessions, from relatively robust accessions such as Mt Eitan (B1K-49 site; mean delta of period (dPeriod) of 0.39 h±1.0) to highly plastic ones as Talkid stream (B1K-08 site ; dPeriod of -4.73 h±1.2) (Table S1). Changes in the amplitude of the clock were even more dramatic between the two environments with mean values doubled under HT compared to OT (Fig. 1b; Table S1). As with period, the plasticity in the amplitude also varied between increases of 10% (B1K-37) to more than 400% (B1K-44; Table S1); yet overall there was no overlap between amplitude levels under two environments. Notably, this clock plasticity results are very much alike to those weobserved for the bi-parental wild barley doubled haploid population (8).

**Figure 1:**
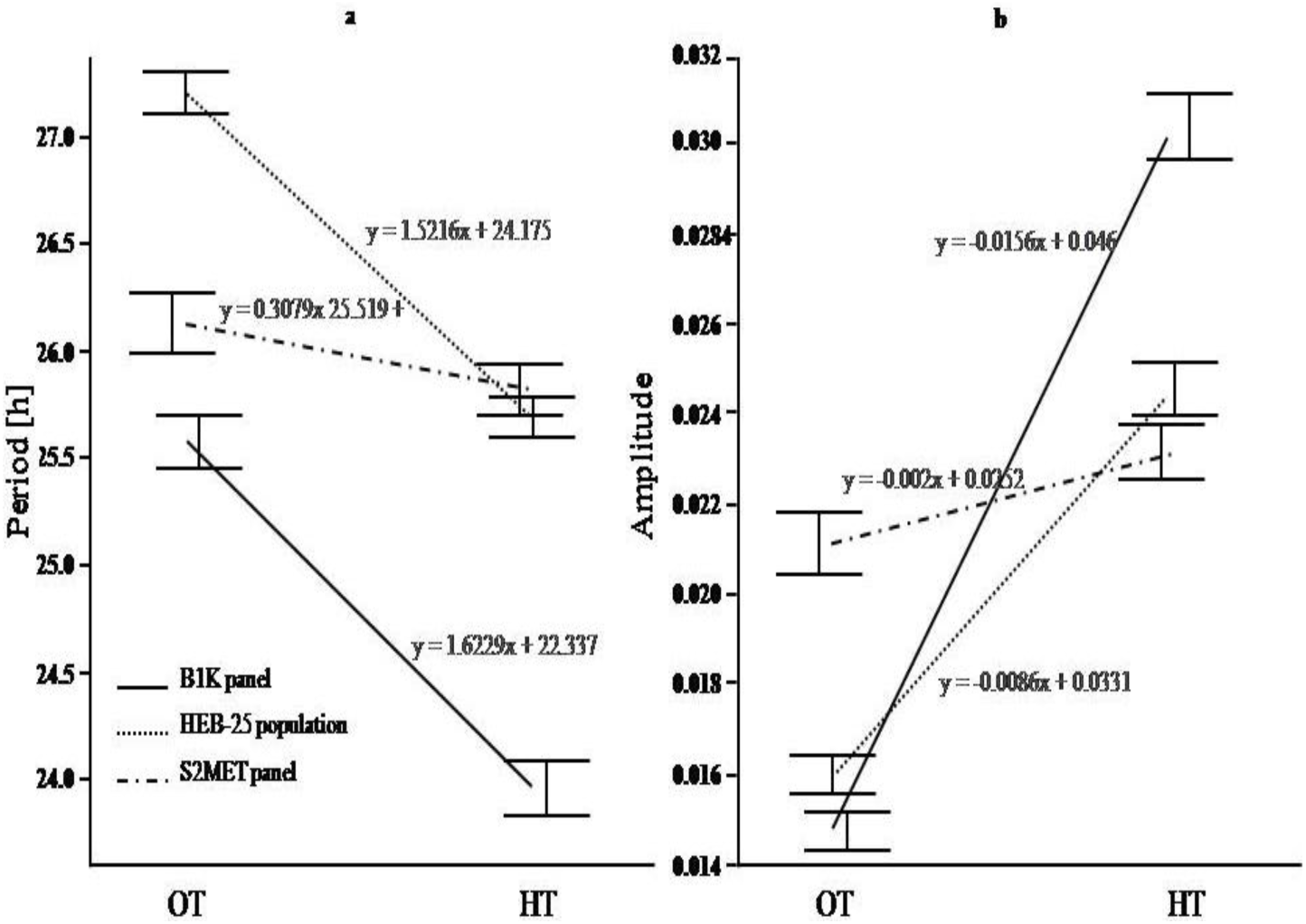
Differential rhythms and thermal plasticity between barley wild panel (Barley1K, B1K), interspecific multiparent population (HEB) and cultivated breeding (S2MET) plant material. Circadian rhythms reaction norms of barley wild *H. spontaneum* panel (n=283), HEB-25 interspecific multiparent population (N=338) and advanced cultivated *H. vulgare* breeding material (N=232) under two thermal environments, optimal temperature (OT; 22°C) and high temperature (HT; 32°C) for clock period (A) and amplitude (B).

Next, we analyzed the interspecific multiparent barley population HEB-25 that includes 25 wild barley donor accessions into the background of the cultivar Barke (26). In this population, each of the lines is theoretically homozygous for the cultivated genome at 71.875% of the sites (26), as compared to none such cultivated diversity in the B1K. Nevertheless, each HEB family is derived from a different *H. spontaneum* accession that originate from the wide barley distribution range including Tibet, Fertile Crescent, and Southern Levant (26). Thermal plasticity of the clock period was similar in direction to that found in the B1K, i.e. a mean dPeriod of -1.52 h± 2.53; Fig. 1a; Table S1). However, we observed a much less significant difference in the changes of the clock amplitude between the HEB and B1Kcollections (Fig. 1b); while the dAmplitude in B1K averaged at +142.6%, that in the HEB lines was half of that (dAmplitude= +72.3%). Since the HEB population is composed of 25 sub-population we partitioned these comparisons to each of the families. This analysis showed that, as between the B1K sites, there was a spectrum of responses between the families with different HID (*H. spontaneum* accessions) donor (Supplementary Fig. S1). For example, while the dAmplitude in the HEB-02 (wild donor is HID004; (26)) population was on average +180%, that of lines derived from of HEB-14 (HID144) was almost unchanged (dAmplitude=-4.2%; Supplementary Fig. S1 and S2).

Finally, we analyzed the circadian clock responses of elite barley breeding material for the same environmental changes. The US Spring Two-Row Multi-Environment Trial (S2MET) panel is including232 breeding lines (27), and to the best of our knowledge there was no wild barley material included in its development. A clear difference between the wild and cultivated plant material is the deceleration of the rhythm and increase in the amplitude under optimal temperature in the cultivated barley(Fig. 1a and 1b). While the mean of the wild B1K accessions under OT averaged at 25.5 hr, ranging between 21.04 and 33.1 hr (Table S1), that of the US panel showed a decelerated pace with an average period of 26.1hr under OT. The amplitude of the cultivated material was significantly higher than the wild accessions, showing on average increase of 131% (Table S1; Fig. 1b). These results show clearly that under domestication, as observed for tomato(28), the circadian clock has decelerated, and moreover, under optimal conditions the amplitude has significantly increased. Moreover, unlike the clear plasticity observed in the fully wild (B1K) and interspecific (HEB) populations (Student’s t-test; P<0.0001 different than zero), the mean plasticity of the *H. vulgare* S2MET lines was not significantly different than zero for period (Student’s t-test; P =0.1) and small difference observed for amplitude (Student’s t-test; P=0.01). These differences between the three gene pools are clearly viewed in the reaction norms between OT and HT (Fig. 1a and 1b). Moreover, analysis for the interactions between the gene pools and the environments support these significant differential responses (ANOVA test; P<0.0001) between cultivated, semi-cultivated and wild gene pools (Table S2).

### Genome wide association study(GWAS) identify genetic network betweenDrivers of Clock (DOC) loci in interspecific population

To identify wild alleles that contribute to these changes in pace and plasticity of the clock between wild and cultivated gene pools we performed a genome scan using clock phenotypes and SNP data available for the HEB-25 population (29). Because the choice of the method used to map the trait plasticity has significant consequence on the results (30), we used the circadian clock phenotype dataset to specify these loci by using three different genetic models (see Methods). Our assumption is that signals that will be repeated in two of the three methods (LMM, MLM and EBL) methods are more reliable and deserve follow-up analysis. We also added to the single point analysis a two-dimensional genome scan to capture possible di-genic epistatic interactions underlying the plasticity.

After filtering the SNP data, based on maximum missing genotypes per marker of 25%, and minor allele frequency above 3%, we continued the analysis with 3,013 loci (Table S3). Genome scan for the loci affecting trait per-se (under OT or HT), or their plasticity (delta of trait), was initially conducted with a linear mixed model (LMM), which took into consideration the population structure and HEB familial relationships (see Methods). By using a significant threshold determined by Bonferroni correction or BH FDR 0.1 we identified only three significant QTL. One, which we named Driver of Clock 3.1(*DOC3.1*), resides on chromosome 3 (position 29,085,440 -36,987,723; PVE= 4.5 %) and was associated with variation of the amplitude only under HT(Fig. 2a; Table S4a and S4b). Another significant QTL loci, *DOC3.2* (43,840,769-51,509,488) resides on chromosome 3 nearby to the *DOC3.1*and the most significant loci in this region is BOPA2_12_31475 (position, 51,509,488, LOD = 4.63; PVE= 4.5 %). Notably, the increase in the amplitude under HT for both*DOC3.1* and *3.2* loci was attributed due to the effect of the wild allele for all the markers in these loci (Table S4b, S4c). The third QTL loci named as *DOC5.1*, resides on chromosome 5 (position 605,805,151-608,879,935; and was associated with variation of the period only under OT and the significant marker associated is SCRI_RS_196175(607,080,381, LOD = 4.94 & PVE= 9.8%)(Fig. 2b; Tables S4a and S4b). Also, the shortening of the period by this locus is associated to the wild allele effect for allthe markers in this region (Table S4b, S4c). Figure 2c and 2d depict the marker effects of *DOC3.1* and *DOC5.1* on amplitude and period plasticity. While *DOC3.1*is inherited with dominancy to the cultivated allele (Fig. 2c), *DOC5.1* seemed to be inherited with dominancy to the wild allele (Fig. 2d). It is important to note that onlythe *DOC3.1, DOC3.2*&*DOC5.1* consistently appeared in three of the methods of GWAS i.e LMM, EBL and MLM (Supplementary figure S3).Apart from these, other DOCs appear for the HEB panel such as *DOC5.2* and *DOC6.1* (Table S4), however they are the results from the Tassel MLM model only with a permissive p-values.

**Figure 2:**
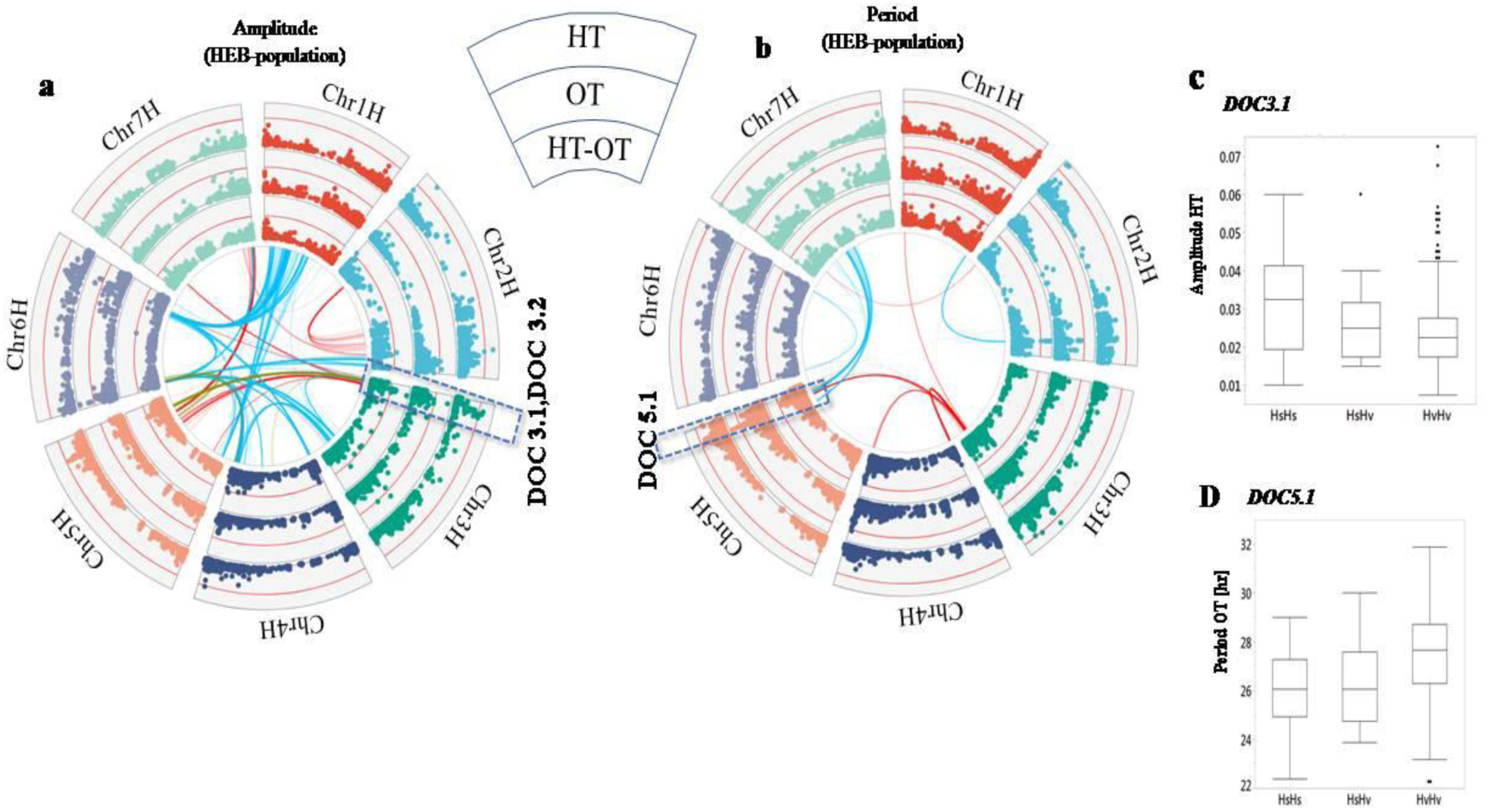
Circos plots depicting the GWAS results for (a)Amplitude and (b)Period of the clock in HEB population.Barley chromosomes in the plot are depicted in different colors.Outer, middle and inner Manhattan plots indicate –log_10_(*p*) of one-dimensional GWAS for high temperature (HT), optimal temperature (OT) and for the delta (HT-OT), respectively. Red lines in the Manhattan plots indicate significant threshold (*p* = 0.001). Links in the center of the circles indicate significant di-genic interactions detected by two-dimensional two-locus genome-wide association study (*p* = 0.001). Red, blue and yellow links indicate high temperature, optimal temperature and the delta, respectively. *DOC3.1, 3.2*&*5.1* appear in all the three methods (LMM,MLM and EBL) analysis and hence are demarked in the circos. Boxplots of clock plasticity for (c) *DOC3.1*effects on Amplitude and (d) *DOC5.1*effects on Period. HsHs, homozygous for wild allele; HvHs, heterozygous for wild and cultivated allele, andHvHv, homozygous for cultivated allele.

Next, we wished to expand our analysis and look for possible epistatic interactions that may explain the two type of changes in the clock (deceleration under domestication and thermal plasticity). We performed a two-dimensional(2D) GWAS with the results obtained with the HEB population under OT and HT (see Methods). Table S5 summarizes all the significant di-genic interaction that we identified for the different clock traits. Some of the loci that played a role in these interactions were not found in the previous 1D GWAS. We noticed that the interactions between the loci appearing with high additive values are mostly heat-conditioned and act on clock amplitude(Table S5; Fig. 2a; Supplementary Fig. S4). At the same time, conditioning is also found across period (Supplementary Fig. S5) and amplitude, where each member of an interacting pair condition a different trait for the other. For clock amplitude under HT we identified two major interactions between *DOC3*.1 or *DOC3.2*, which we identified in 1D analysis, with newly identified loci at of chromosome 5 (Table S5).Figure 3 is depicts these interactions for both *DOC3.1* and *DOC3.2* with the same locus on chromosome 5. While the interactions of *DOC3.2* were more significant for amplitude and under HT only (Fig. 3a and 3b), those of *DOC3.1* acted reciprocally between the chromosome 3 and 5 on both amplitude and period (Fig. 3c and 3d). This HT-dependent interaction showed that the modulation of clock by *DOC3.1* is conditioned by the allelic combination in chromosome 5 and vice versa. The increase of the amplitude between homozygous for wild and cultivated allele at *DOC3*.1 (*DOC3.1*^Hs/Hs^ vs *DOC3.1*^Hv/Hv^) is conditioned by homozygosity for the wild at the chromosome 5 locus (Fig. 3c). Similarly, but this time for the period, the acceleration of the clock by the chromosome 5 QTL is conditioned by homozygosity at the *DOC3.1* (Fig. 3d).To summarize, these clear digenic interaction (with one locus not identified in1D GWAS) provides amplitude and period plasticity only for the carriers of the wild alleles at both loci.

**Figure 3:**
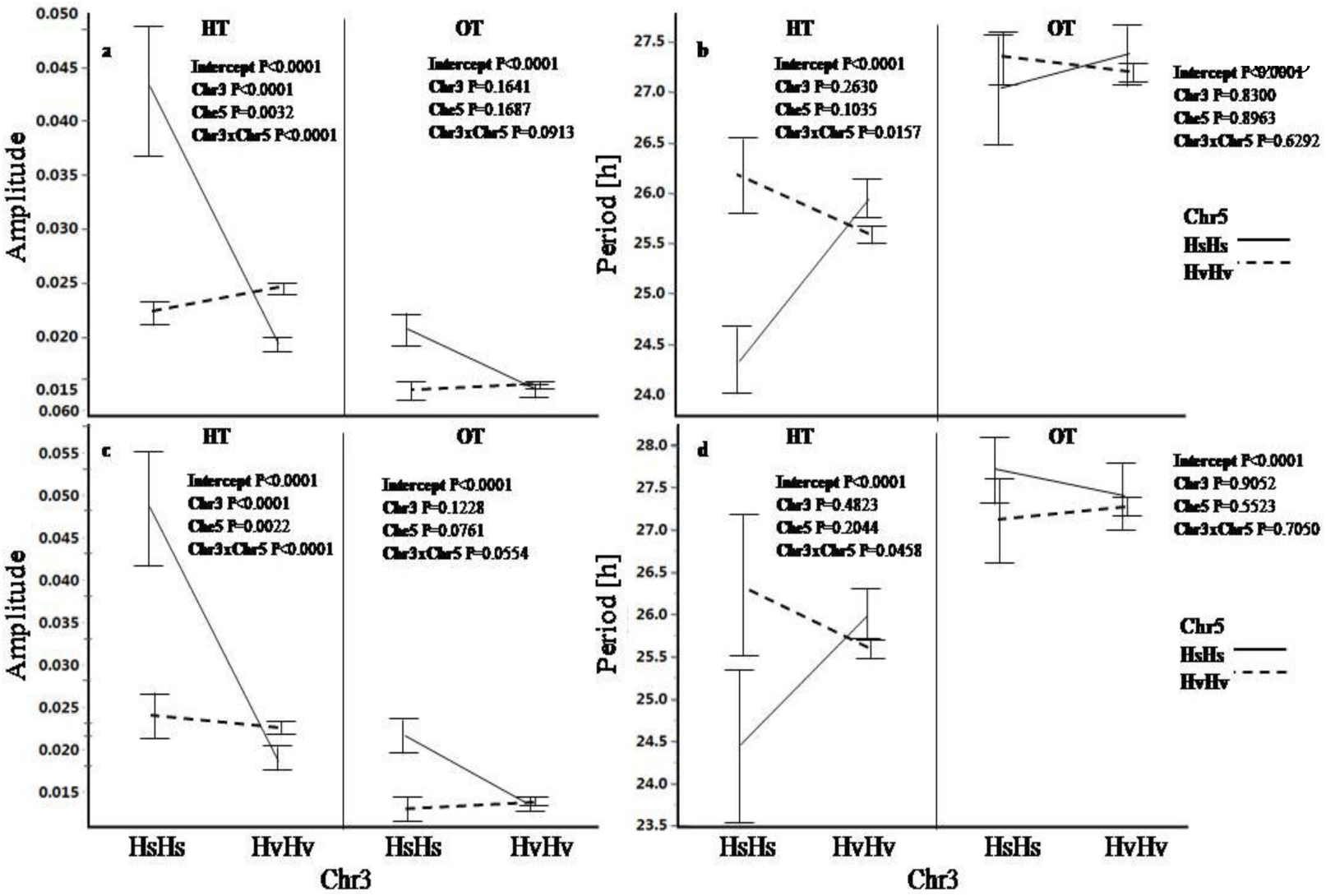
Epistatic interactions between DOC loci on chromosome 3 and 5 for clock amplitude and period.Least-square mean value comparisons (reaction norms) for the (a and b) *DOC3.1*(position 30,927,745) and (c and d) *DOC3.2* (position 47,570,630) genotypes under optimal temperature (OT; 22°C) and high temperature (HT; 32°C) in the HEBpopulation.A two-way ANOVA including each pair of lociin chromosome 3 and 5 tested if the interaction is significant.

### GWAS for clock diversity in breeding material highlight differential genetic network and lack of epistatic interactions

To unravel the genetic network underlying variation in the clock phenotypes, including that underlying thermal plasticity, we repeated the two types of GWAS, i.e. single point and di-genic (1D and 2D), for the S2MET breeding panel using the SensyPAM data we obtained under OT and HT (Table S1). Overall, this genomic architecture included a different set of significant QTL that we associated with clock period, amplitude and their thermal plasticity (Fig. 4). Interestingly, for this breeding panel we did not find any QTL that came consistent with the different models we used. From Tassel MLM,we identified onelocus underlying amplitude variation on the telomeric end of chromosome 3 on the, i.e. *DOC3.4*, explaining 13.9% and 12.5 % of the amplitude plasticity(delta Ampitude) and its variation under OT, respectively(Fig. 4a, Table S4; see Methods). The most notable locus associated with variation in amplitude of the clock, *DOC7.2b*, explained 14% of the variation for the trait under optimal thermal condition. Finally, we associated one additional QTL, *DOC2.2b*, affecting variation under high temperature on chromosome on chromosome 2. We noticed that loci underlying the period variation, as these affecting amplitudes, in this US panel had higher contribution to the variation. This included *DOC3.3* that explained 10.7% of general period variation under HT, and *DOC2.3* and *DOC5.3* with PVE=8.3% and 8.7%, each is higher from any locus in the HEB population (Table S4).

**Figure 4:**
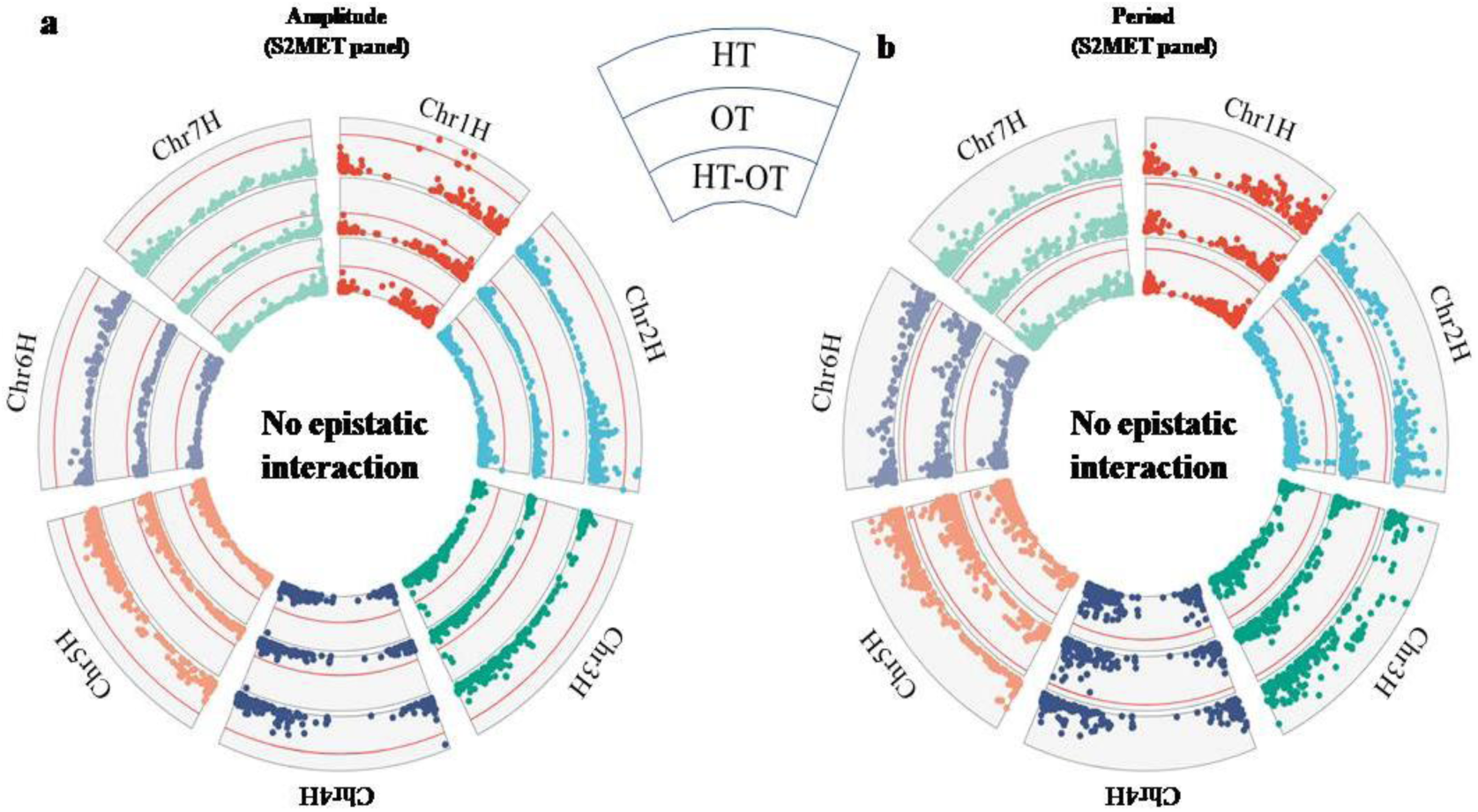
Circos plots depicting the GWAS results for (a) Amplitude and (b) Period in S2MET panel.Barley chromosomes in the plot are depicted in different colors. Outer, middle and inner Manhattan plots indicate –log_10_(*p*) of one-dimensional GWAS for high temperature (HT), optimal temperature (OT) and the delta between values (HT-OT), respectively. Red lines in the Manhattan plots indicate significant threshold (*p* = 0.001).

In addition to the interspecific and cultivated populations identifying a different set of DOC QTL(Fig. 2 vs Fig 4; Table S4), the cultivated population exhibited a lack of significant epistatic interactions was, under both temperatures. In any method taken (see Methods) we could not identify in the breeding panel significant di-genic interactions for any of the clock traits (Supplementary Fig. S6 and S7). This is compared to ample amount of significant interactions found for trait per se (amplitude and period) and their plasticity (dAmplitude and dPeriod) in the HEB population (Table S5; Fig. 2).

### The DOC3.1 and DOC5.1are significantly associated with temperature-dependent effects on heading and grain yield

Since we observed a significant loss of plasticity from the wild to the cultivated gene pool and no overlap in loci detected for clock variation between the two populations, we were curious to test if some of the DOC loci identified in HEB population could be associated with other traits in cultivated barley.. The S2MET panel was phenotyped for 14 important traits in 39 location-year environments between 2015 and 2017 (27).This allowed us to perform per-trialGWAS for two main fitness traits, i.e. heading date (HEA) and grain yield (GY), using the 5068 high-quantity filtered SNPs (Table S3 and S6). Then, we compared the chromosomal location of the DOC loci identified in the HEB or US panel and checked for co-localization of clock and agronomic trait QTLs. Notably, in these GWAS analyses, SNP associated with heading datewere much more consistent across the 39 environments compared with those associated with grain yield implying heading time QTL are more stable than yield. For example, marker S2_ 429692133 on chromosome 2 (physical position 429,692,133) appeared to be significantly linked(LOD>3) with heading date in 16 out of 39 (41.02 %) field trials (Table S6). Other heading date markers, that were reproducible across different sites ranged from two to twelve environments whereas the upper limit for yield was (7.6%). To represent the frequency of such stable SNP across different sites, we considered the number of significant SNP and generated a density plot for GY and HEA(Fig. 5a and 5b). In that regard, *DOC3.1* and *DOC5.1* belongs to be more as relatively stable loci for GY, i.e. we identified *DOC3.1*significanteffects on GY in up to 3 experimental trials(7%), and for *DOC5.1*, the effect was repeated up to 2 trials depending on the markers we used in the genetic analysis (5.1%; Fig. 5a, Table S7). We could also observe that *DOC5.1*, unlike *DOC3.1*, was also having relatively consistent effect on HEA up to 3 trials(7%) for markers used in the interval (Fig. 5b, Table S6).

**Figure 5:**
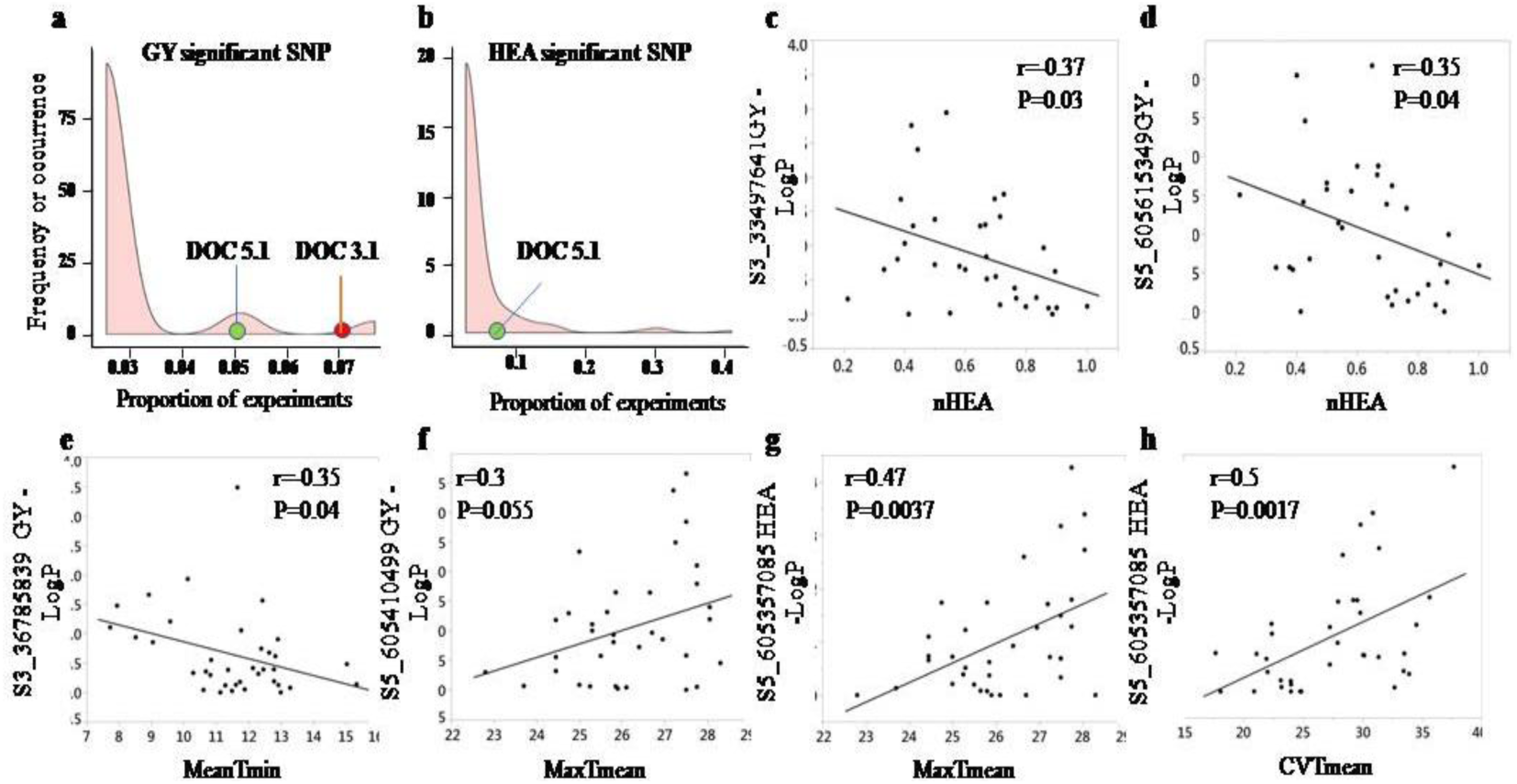
QTL-environment association for DOC loci show their stable and temperature-dependent effects on heading date (HEA) and grain yield (GY) variation in S2MET panel grown across US.Density plots representing the frequency SNP that aresignificantly (LOD>3) associated with (a) HEA or (b) GY variation. Frequency or occurrence (X-axis) is expressed as proportion of experiments (out of 39) in which association marker-traits is significant.Red and green dotson the X-axis demarcate the occurrence of *DOC3.1* and *DOC5.1* loci for GY and HEA. The correlations between normalized heading date at experimental site and significance of the (c)*DOC3*.1 or (d)*DOC5.1* association with GY in the S2MET panel. The correlations between Tmin in experimental sites and GWAS –log_10_(*p*) values for (e) *DOC3.1* and (f) *DOC5.1* for GY. *DOC5.1*-temperature association is depicted by the positive correlations of (g)minimal temperature (Tmin) and (h)CV of Tminwith GWAS –log_10_(*p*) values for *DOC5.1*effect on GY in S2MET panel.

Moreover, we wished to take a more quantitative approach and examine if the relative level ofgenetic association, i.e. its significance, with GY and HEA are conditioned by the environmental variation, therefore suggesting that they are involved in local adaptation.To test the possible relationship between environmental gradients and effect of the DOC loci, we correlated the per-trial GWAS –log_10_(*p*) values of the two traits (HEA and GY) for*DOC3.1* and *DOC5.1*with the environmental covariates (e.g. temperature and rainfall) recorded during these experiments. Also, we made these correlations between GWAS –log_10_(*p*) values and the calculated variation of these environmental covariates, e.g. the coefficient of variation (CV) for minimal daily (night) temperature (Tmin) that represent the stability of the environmental conditions (see Methods). Finally, we made these correlations with regard to the mean normalized flowering time in each location to askif the effect of the DOC locus is more or less effective depending on the stabilityof heading. The full details of the correlation coefficient between the per-trial lGWAS –log_10_(*p*) values and the different environmental covariates are depicted in Table S8. Generally, we found a significant negative correlation between –log_10_(*p*) values of *DOC3.1*markers for grain yield and the mean normalized headingdateof the whole panel on trial(r = -0.37, P=0.03; Fig. 5c). We found similar correlation between HEA stabilityof the trial and the significance of the *DOC5.1*effects on grain yield (r = -0.35, P=0.04; Fig. 5c). These results suggest that the more variable was the flowering between lines the higher was the effect of the *DOC3.1*or *DOC5.1* on grain GY variation.

When we tested the correlations between the environmental covariates and the effect of the two loci on GY or HEA we found a difference between the two loci. For *DOC3.1*, we could identify a significant negative correlation (r=-0.35, P=0.04) between Tmin in the site of experiment and the specific *DOC3.1* GWAS –log_10_(*p*) for GY (Fig. 5e). On the other hand, the correlation between GWAS -Log10(P) of GYwith temperature or other environmental gradients in sites were marginal. For example, the highest correlation of r=0.3 (P=0.055) was observed between the range of maximal daily temperature (maxTmean) and –log_10_(*p*) of GWAS for GY (Table S8; Fig. 5f). Nevertheless, the correlations between environmental covariates in experimental site with the GWAS –log_10_(*p*) values for *DOC5.1*on HEA were much more significant and abundant. Moreover, these correlations appear both for the means per se, as well as for the CV of the temperatures and precipitation values. We found a strong positive correlation between per-site –log_10_(*p*) for HEA with Tmax (r=0.47, P=0.0037; Fig. 5g), as well as significant correlation with CV Tmax(r=0.5, P=0.0017; Fig. 5h). These results indicate higher and/or unstable temperatures during growth periodare associated with the effect of the *DOC5.1* on the flowering time variation.

Finally, we made these quantitative correlations between environmental covariates and genetic significance for the DOC loci identified in the US S2MET panel (Fig. 4; Table S4a). It is interesting to note that unlike the environmental-QTLcorrelation for effects on GY we identified for the wild allele of *DOC3.1*, we could not find any such relationship (P>0.05) for any of the DOC loci identified in the GWAS of this material. These QTL-environment correlations could only be found for the effects of*DOC3.4* and *DOC5.3*on HEA variation (Table S8).

## Discussion

### The nature of circadian clock changes under domestication

A clear difference between wild and modern breeding material is that wild accessions accelerate their rhythmicity and increase their amplitude oscillation under heat, while the cultivated genotypes are much less responsive and overall maintain a similar peripheral clock (Fig. 1). In the current study we used a high throughput readout of non-photochemical quenching (NPQ) to obtain measures of clock rhythmicity (period and amplitude). Previously, Dakhiya et al. (7) showed that NPQ oscillation gives a fairly good proximation for the clock rhythms by comparing clock genes to F-based readouts in Fytoscope, mainly by using Arabidopsis mutants, i.e. *cca1* and *lhy*. Nevertheless, we showed that genetic changes in these rhythms are not always correspondent to the responses of the core clock genes, e.g. CCA1, and that these probably correspond better to peripheral rhythms such as photosynthesis. For example, the effect of the wild barley allele at *frp2.2* locus accelerated the clock under heat yet examination of CCA1 and TOC1 temporal expression did not find significant effect of this locus on this core clock gene (8). Nevertheless, despite the fact that as wild barley was faced with a more extreme environment compared tocontemporary breeding material, there seems to be an advantage for plasticity of these rhythms, either directly affected by the core clock or in its transduction to peripheral activities. Accelerating the clock suggest that in a similar manner to seasonal escape mechanism, where cereal that flower earlier avoid detrimental conditions for growth and reproduction phase(31), higher daily pace may support avoidance of physiological activities in more stressful periods of the day.. Further detailed physiological and molecular analysis of nearly-isogenic lines for the DOC identified in this study, including the differential transcriptome and metabolome changes, should unravel the main pathways that will explain this accelerating strategy and its possible benefits for fitness.

### Genetic diversity underlying loss of clock plasticity under domestication

The type of genetic material investigated in this study, as well as other comparisons between genotypic and phenotypic diversity under domestication(32–34), consider nuclear genome diversity that had obviously undergone a bottleneck (35). Moreover, recent studies are attempting to identify signatures of selection by identifying genetic sweeps across these nuclear genomes between wild and cultivated panels (36, 37). Interestingly, few of the loci we identified in this study overlap with some of the resequenced genes reported under selective sweep between wild and cultivated barley (38). This is including overlapping of the *DOC3.2* loci marker SCRI_RS_141171 (43,840,769) with that of BOPA2_12_30924, which is located on chromosome 3 631,804,839. Pankin et al.(38)found that the gene HORVU3Hr1G090440.4 spanning 631,804,086-631,808069 is under selection. However, since at least one of the major DOC loci we identified in this study, *DOC3.1*, has not been included in the list of candidate domestication genes, it is still an open question as to loss of these DOC loci resulted in a a signature of selection. Combining our top-down QTL approach with such bottom-up analysis could be instrumental in finding the causal variation underlying functionality of genes and their selective advantage.

Furthermore, with regard to the circadian clock variation and its change under domestication, as we report in this study (Fig. 1), there may be additional overlooked source of diversity that went through a genetic sweep and which has affected the loss of plasticity in the cultivated breeding material. Previously, our study of reciprocal doubled haploids within wild barley have shown a significant difference in the thermal response of the circadian clock that we could link with plasmotype diversity (8). This variation corresponded to 6 non-synonymous mutations between the founders of the populations. This suggests that in the search of variation on the clock, we should consider and design experiments that will examine the direct links between plasmotype diversity and circadian clock variation under different environments, as well as cytonuclear interactions. Such designs should also consider the fact that there is a biologically relevant interaction between the chloroplast and nuclear genomes, i.e. retrograde signaling(39), and that effects of nuclear genes on the clock and its output could be conditioned by protein or RNA partners encoded in the organelle. In fact, we recently generated a relevant cytoplasmic multi parent population (CMPP) that includes introduction of ten wild barley cytoplasm into the background of the cultivated barley(40). This new fully-homozygous infrastructure will allow us to explore these cytonuclear interactions for different phenotypic layers including circadian clock and agronomic traits in multiple sites, and to investigate in depth how such pleiotropy is maintained or lost in accordance with environmental changes.

### QTL-environment association for detecting local adaptation

Recently, Wadgymar et al. (41) proposed a framework for evaluating the nature of local adaptation including distinguishing between genetic tradeoffs and conditional neutrality of QTL involved in local adaptation. They, and others (42) proposedthat loci that have effects on fitness in one environment, but not in alternative environments (i.e., conditional neutrality), appear to be more common. From an agricultural point of view, and obviously for the same reason of local adaptation, crop breeding is done in the target area to allow better relevance of the material. Indeed, same group took this approach and tested a panel of switchgrass and showed that adaptive trait variation has beneficial effects in some geographic regions while conferring little or no detectable cost in other parts of the geographic range (43). Similarly, herewe demonstrate that among the significantly associated loci the percentage of markers repeated in more than several sites/years is low for GY and much more for HEA (Fig. 5a and 5b). This could be because GY is a complextrait with relatively low heritability that influence from several other traits(44). Nevertheless, the markers found for clock rhythmicity (DOCs) are found significant for HEA and GY in more trials than other markers, suggesting that they are involved in local adaptation. Moreover, we also provide an alternative quantitative approach to test the level of local adaptation with regard to the environmental agents involved. This is simply done by performing per-trial GWAS to obtain the significance (P value) of the SNPs and then perform correlation, or other similar association test, to examine relationship of significance with environmental co-variates. This approach is different than counting the frequency of single nucleotide polymorphisms (SNPs) associated with fitness and how it co-varied with climate across the range of experiments conducted (Fournier-Level et al. 2013).

### The adaptive value of plasticity vs robustness under domestication and its molecular basis

In a previous study, we showed that within the wild barley populations there is a standing variation for the circadian clock plasticity and that that it varies between accessions adapted to different niches. For example, the B1K-09-07 that served as a parent of the ASHER DH population showed significant shortening of the clock rhythm (by more than 3 hr; (8)). This is compared to relative robust clock of B1K-50-04, an accession from the colder Northern part of the B1K collection (25), for which the period shortened slightly by one hour. This, and the fact that a significant relationship between temperature at the site of collection and the clock period was found indicated the adaptive value of the clock to changing environment in the wild(7). But how would that be relevant in agriculture, and why do we observe loss of such plasticity, as exemplified by overall lesser response for both period and amplitude (Fig. 1), and also by loss of significant plasticity QTL such as *DOC3.1*? One obvious reason might be that modern cultivars were bred in a more stable environment than their wild ancestors, and more specifically, more homogeneity in sites in the different sites of cultivation vs growth of wild populationsin wide adaptive niche (25).

Vis-à-vis adaptation under domestication for barley and lack of environmental responsiveness involved, it was shown previously that misexpression of barley ortholog for the Arabidopsis circadian-clock gene *ELF3* of circadian clock gene in the *eam* mutants (19). This mutation allowed early-flowering day-neutral phenotype with rapid flowering under either short or long day (45). This may suggest that under cultivation loss or change of circadian clock functions such as light responses have been favored, e.g. deletion and mutations in the EID and LNK2genes in cultivated tomato found to be responsible for the clock deceleration(24). Since a direct link between allelic changes from wild to the cultivated tomato for yield has not been shown yet, but only that of growth (24), it is still remains to be studied what is the adaptive value of these mutations. Besides, are loss of function for clock genes is typical to these changes under domestication as reported so far? and if not, and functional alleles were retained, could that suggest a repurposing of the genes for rhythmicity, and its plasticity, to yield adaptation? Further isolation of the causal genes underlying DOC and the study of their possible pleiotropic effects on agronomic traits, as well their biochemical functions, will be required to answers to these questions.

## Materials and methods

### Plant material and genotypic data

For this clock thermal plasticity analysis, we used tree Barley panel: wild barley (*Hordeumspontaneum*), interspecific multiparent barley population HEB-25 and breeding cultivated panel (*H.vulgare*). We selected two hundred and eighty-three accessions of wild barley which are single-seed descents from our original Barley1K collection (25). The B1K accessions were collected in hierarchical manner (5 microsites in each location)from 51 sites that represent a broad genetic and ecogeographic adaptation. We initially attempted to select one representative accessionsfrom each micro-site and more from sites showing relatively higher genetic diversity based on SSR analysis of the collection (25). For the HEB population, we selected three hundred thirty-eight lines that represent all 25 families. These 25 HEB families were generated by introduction of different *H.spontaneum* accessions into the background of the cultivated Barke background (26). All original BC_1_S_3_ lines (two generations earlier) and their corresponding parents were genotyped using the barley Infinium iSelect 9K chip (26), consisting of 7864 SNPs (46). The single seed descendedHEB-25 lines we used for the clock measurements are a BC_1_S_3::5_. Two hundred thirty-two cultivated barley (*H. vulgare*) from the US Spring Two-Row Multi-Environment Trial (S2MET) are an advanced cultivated breeding material for adapting into environmental changes (27). This panel consists of one-eighty-three lines from five U.S. breeding programs and 50 are from crosses between some ofthese lines.

### Clock phenotype under optimal and high temperature

To mimic the natural growth conditions of the wild barley population in the Southern Levant, where the original Barley1K infrastructure was collected (25). Plants were grown to the emergence of the fourth leaf under 10h light and 14h dark, at a constant temperature of 20°C. We grew the two other panels up to the emergence of the fourth leaf under 14h light and 10h darktomimic the spring growing of Barke in Europe and in the US. Following this entrainment of the plants for four weeks, we moved them to the high-throughput SensyPAM (SensyTIV, Aviel, Israel) custom-designed to allow F measurements in up to 240 plants for each experiment (see details at (8)). For the clock measurement, F was measured every 2.5 hours, for 3 days, in continuous light. We measured each genotype twice under each temperature, with 4 to 5 plants included in each round. For the clock analysis, NPQlss ((Fm-Fmlss)/Fmlss) was calculated and normalized by the one control line that appeared in each experiment. The circadian clock free running period (FRP), amplitude (AMP) and amplitude error were extracted using the BioDare2 website (https://biodare2.ed.ac.uk). The input data was set to “cubic dtr” and the “MFourFit” was used as the analysis method (47).

### Statistical analysis

We used the JMP version 14.0 statistical package (SAS Institute, Cary, NC, USA) for statistical analyses and for generating reaction norms for the means and standard errors of the different traits. Student’s t-Tests between treatments were conducted per panel using the ‘Fit Y by X’ function. A factorial model was employed for the analysis of variance (ANOVA, table Sx), using ‘Fit model’, with temperature treatment and panel as fixed effects. The density plot was made using MVAPP(48).

### Genome-wide association study (GWAS)

Weconducted genome-wide association to identify trait variations, per-se, under optimal and high temperatures (OT and HT), and to assessdi-genicinteractions(2D-scan). Since the methods of choice (genetic model and statistics) for the genome scan have a major effect on the loci identified we compared between several options to point into most reliable signals that we could support by more than one method:

### Extended Bayesian Lasso (EBL)

EBL (49)is the extension of Bayesian Lasso (50), that separates the regularisation parameter into a shrinkage factor for the overall model sparsity (*δ*^2^) and a shrinkage factor for individual markers 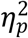. This approach is intended to assign different magnitudes of shrinkage to individual marker effects. In EBL, the following linear model was used:

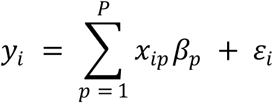

Where *y*_*i*_ is a phenotypic value of individual *i, x*_*ip*_ is a genotype of marker *p* of individual *i, β*_*p*_ is a effect of marker *p*, and *ε*_*i*_ is a residual for the individual *i* with 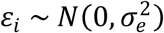. Each regression parameter *β*_*p*_ is assumed to follow

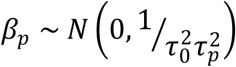

where 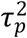 determines the magnitude of shrinkage for *β*_*p*_, and 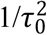 is the residual variance, respectively. Then, 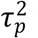 was assumed to follow a prior distribution

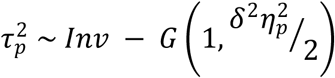

where *Inv* − *G* indicates the inverse Gamma distribution,*δ*^2^ is the shrinkage factor for all markers and 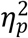 is the shrinkage factor unique to marker *p*. A prior distribution for *δ*^2^ was*δ*^2^ ∼ *G* (1, 1), and for 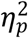 was 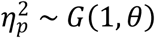 where the rate parameter *θ* is the hyperparameter for EBL. Three values of *θ* were tested: 0.1, 1, and 5and a nested five-fold CV was performed to determine the optimal hyperparameter. The EBL wasperformed by using function *vigor* in the R package “VIGoR” (51). Then absolute value of *β*_*p*_ was used as the GWAS score.

Tassel MLM: Mixed Linear Model(MLM) in Tassel software considers both population structure andkinship in the association analysis. It reduces Type I errordue to relatedness and population structure. MLM was used to identifythe associations between phenotypic and genotypic data in Tassel v5.2.5.0 (52) with optimal compression and variance component estimation by P3D (population parameters previously determined). The P value indicated the degree of associationbetween a SNP marker and a trait, and the R^2^depictsthevariationexplained by the significantly associatedmarkers. Markers with an adjusted -log_10_(P-value) ≥ 3.0 were regarded as significant for all traits.

#### Two-dimensional genome scan (2D-scan)

For2D-scan, we considered following linear mixed models:

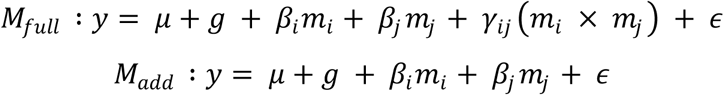

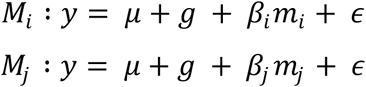

where *y* represents a vector of phenotype,*m*_*x*_ represents coded genotypic values of marker *x*(−1 and 1 for homozygous, 0 for heterozygous), *β*_*x*_ represents effect of marker *x, γ*_*ij*_ represents a vector of coefficients for interaction between marker *i* and *j*(*m*_*i*_×*m*_*j*_), The variable *g*models the genetic background of each line as a random effect with *g*∼ *N*(0, *Kσ*^2^_*G*_), where *K* is a kinship matrix calculated from the nucleotide polymorphisms, and *σ*^2^_*G*_is the genetic variance.*ϵ* represents the residual error such that *ϵ* ∼ *N*(0, *Iσ*^2^_*e*_), where *I* is an identity matrix and *σ*^2^_*e*_is the residual variance.*M*_*i*_ and *M*_*j*_ are equivalent to the model used for single marker based GWAS. *M*_*add*_ is a model to test additive effects of two loci while *M*_*full*_ include both additive and interactive effects of two loci. Then, we derived two *P*-values from the above models.

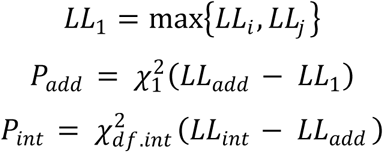

where *LL*_*int*_, *LL*_*add*_, *LL*_*i*_ and *LL*_*j*_ are log likelihood for *M*_int_, *M*_add_, *M*_i_ and *M*_j_, respectively. *P*_*add*_ was used to test significance of addition of a locus to the another. *P*_*int*_ was used to test significance of interactive effect. *df.int* is degree of freedom for two loci interactive effect that is equivalent to number of combination patterns in the given two loci. To solve the linear mixed models used in the 2D-scan, we used the R package “gaston” (53)(54). Maximum likelihood estimates of variance components were obtained using function *lmm.diago*, and log likelihood of each model was calculated using function *lmm.diago.profile.likelihood*.

### Correlations between environmental covariates and GWAS results

The full environmental covariate data appears at Neyhart et al. (27). For the analysis in our current study we calculated the daily mean temperature according to the daily min and max temperature in each field trial. From this data we calculated the mean of the minimal (min), maximal(max) and mean daily temperature, and the range of the mean daily temperature. In addition, we calculated the maximum of the mean and max temperatures and the CV for the min, max and mean daily temperature. We also used the cumulative perception in each trial and calculate the GDD. We also normalized the HEA and GY in each trial following similar normalization conducted by Merchuk et al. (2018) for multiple year

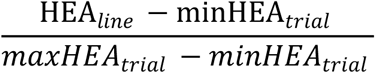

## ACKNOWLEDGEMENTS

This work was supported by the Israeli Science Foundation (ISF) program (1270/17) and the Chief Scientist Competitive Grant (20-01-0080) from the Ministry of Agriculture to E.F.

## Supplementary figures legends

**Figure S1:**
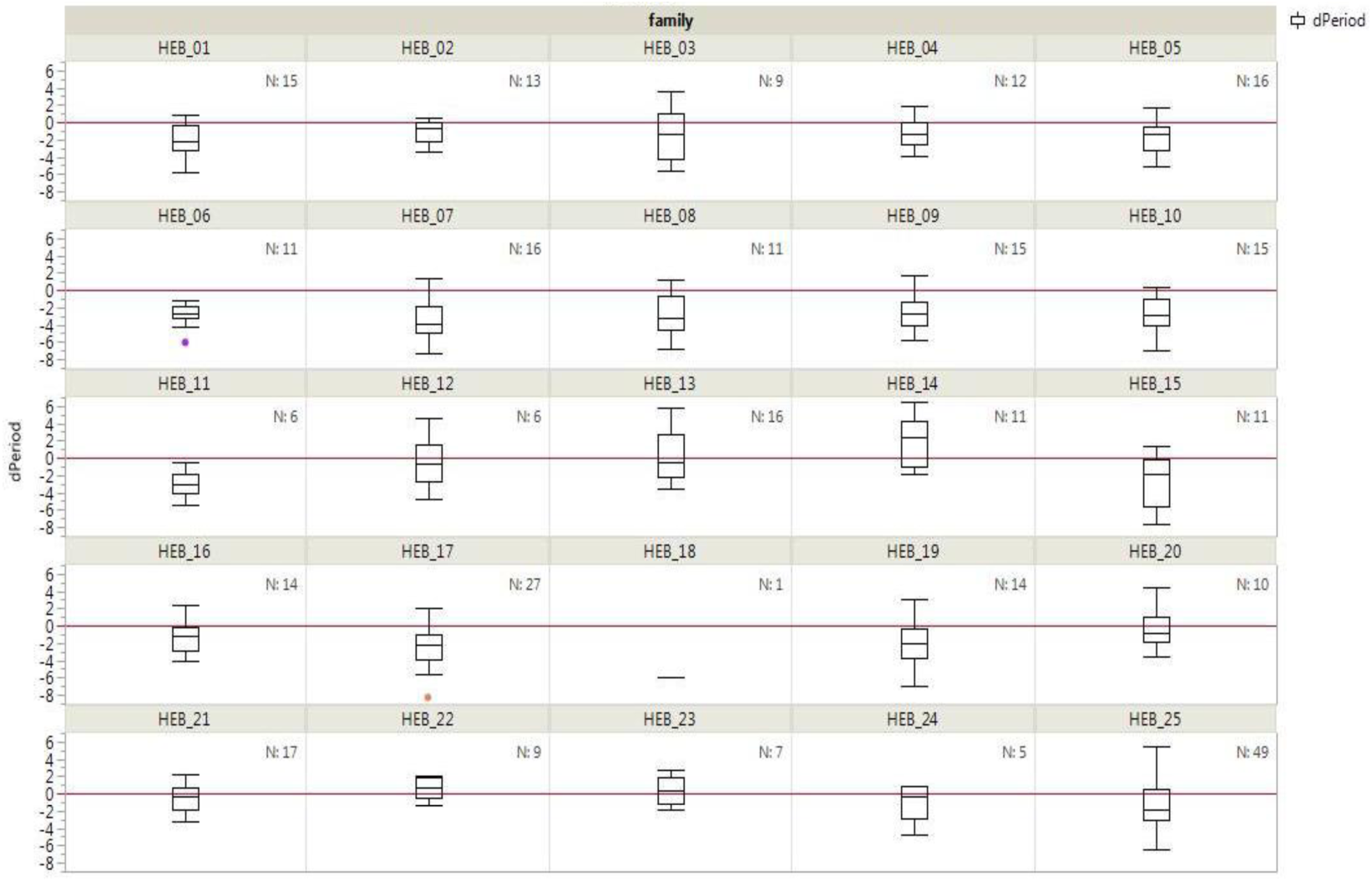
Differential period thermal plasticity in the HEB-25 families. Box plots for dPeriod among each of the sub-populations (each with a different *H. spontaneum* donor; (26)) of the interspecific multi-parent barley population HEB-25. dPeriod is the period under high temperature (HT; 32°C) minus that under optimal temperature (OT; 22°C). The red line represents dPeriod=0 that means no period plasticity.

**Figure S2:**
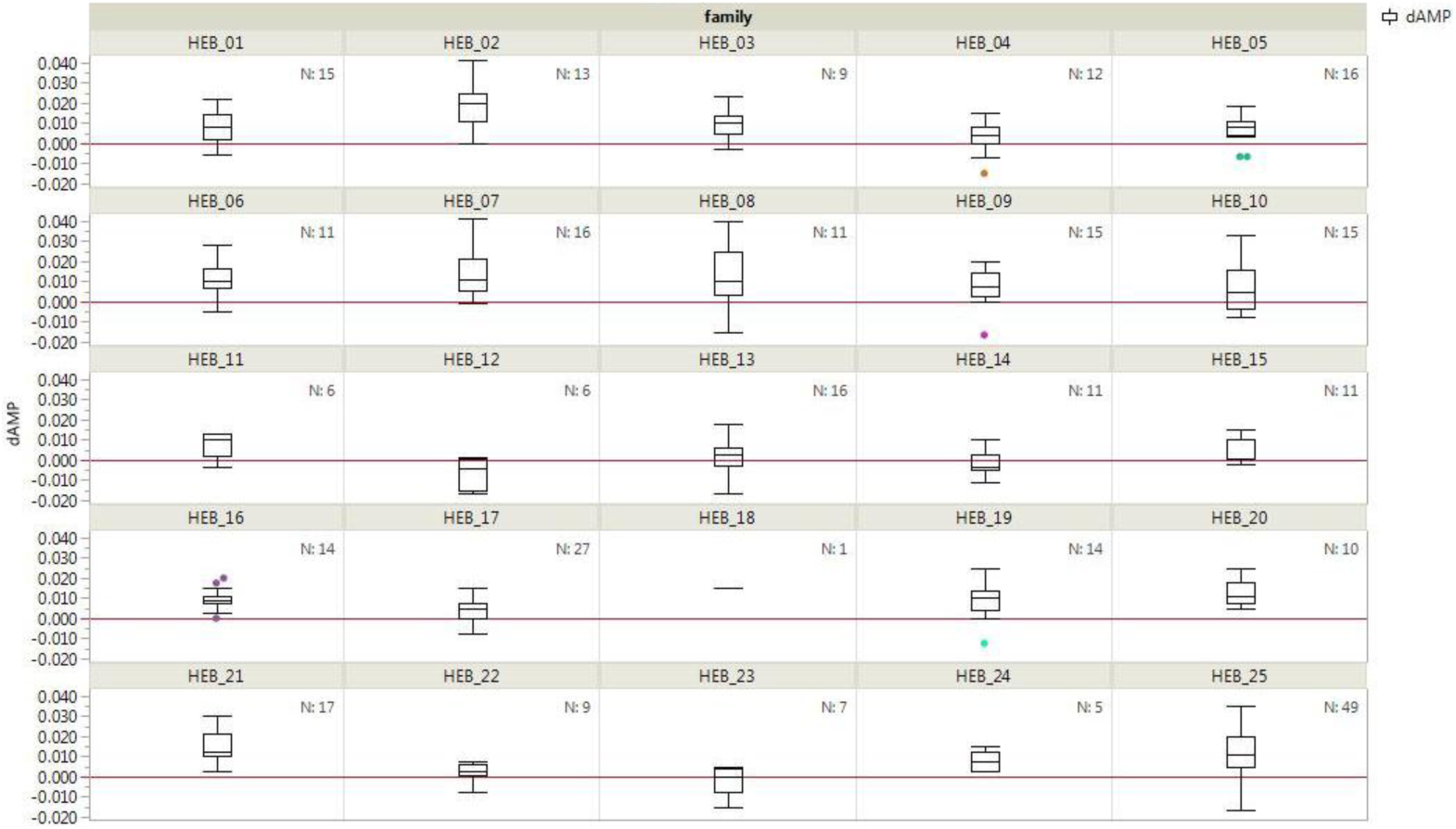
Differential amplitude thermal plasticity in the HEB-25 families. Box plots for dAmplitude among each of the sub-populations (each with a different *H. spontaneum* donor; (26)) of the interspecific multiparent barley population HEB-25. dPAmplitude is the amplitude under high temperature (HT; 32°C) minus that under optimal temperature (OT; 22°C). The red line represents dPAmplitude=0 that means no amplitude plasticity.

**Figure S3:**
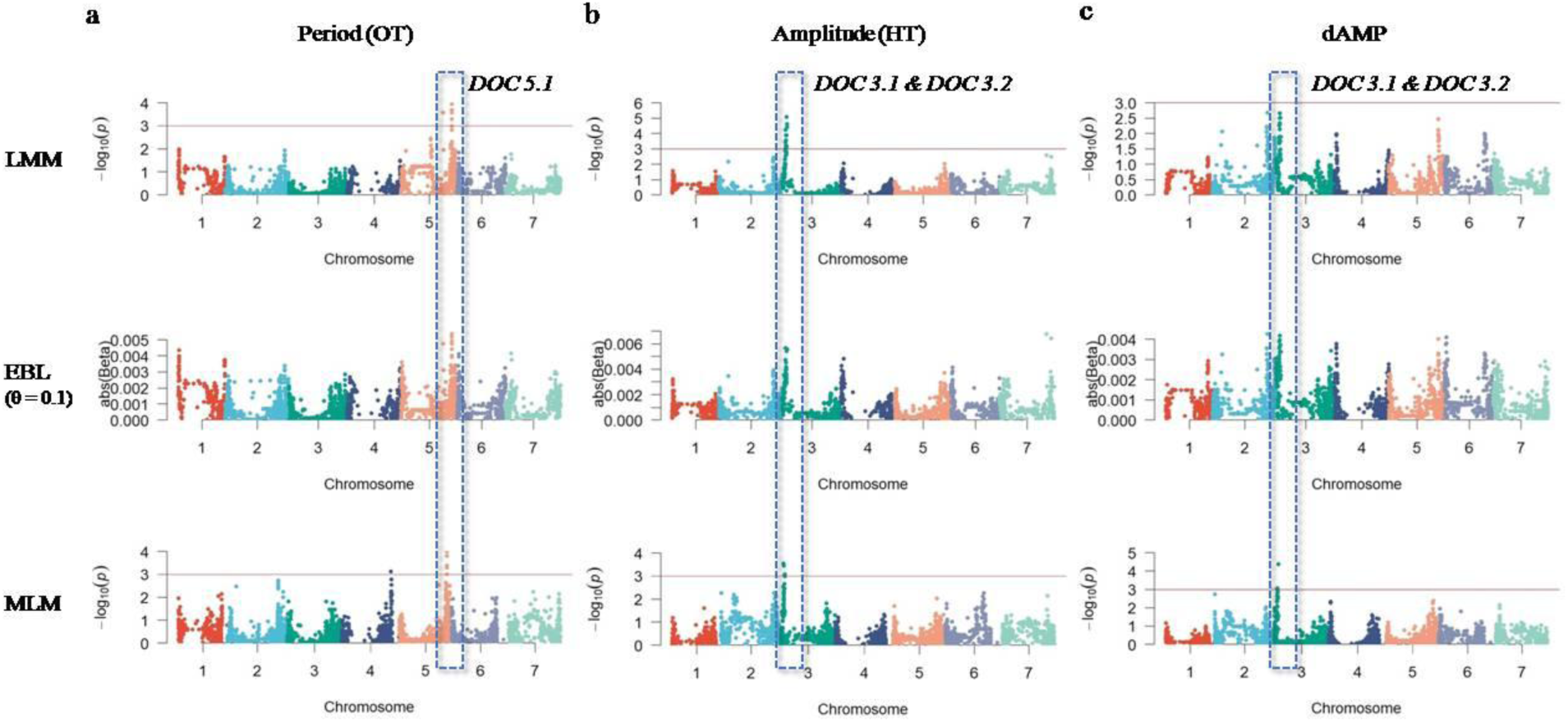
Significant DOCs detected in more than one method (LMM,EBL and MLM) of the GWAS analysis.(a) *DOC5.1*identified for period OT (b) *DOC3.1* and *3.2*identified for amplitude HT and (c)*DOC3*.1 and *3.2* identified for delta amplitude.

**Figure S4:**
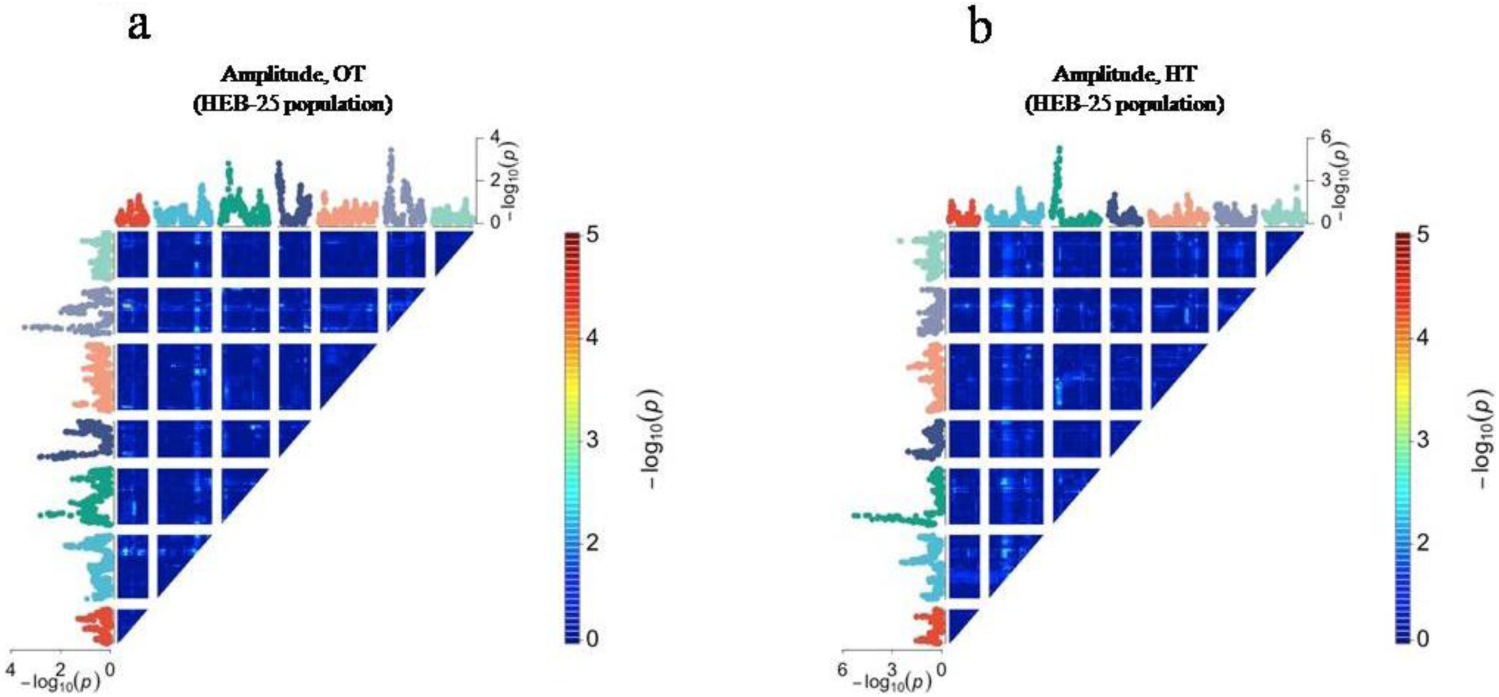
Genome-wide scan for amplitude in HEB-25 population under (a) optimal temperature (OT) and (b) high temperature (HT).Heat map for two-dimensional genome scan with a two-loci interaction model. The Manhattan plots on x- and y-axes in each panel indicate result of one-locus model genome scan. Genome scans were performed using linear mixed model that account population structure as the covariates.

**Figure S5:**
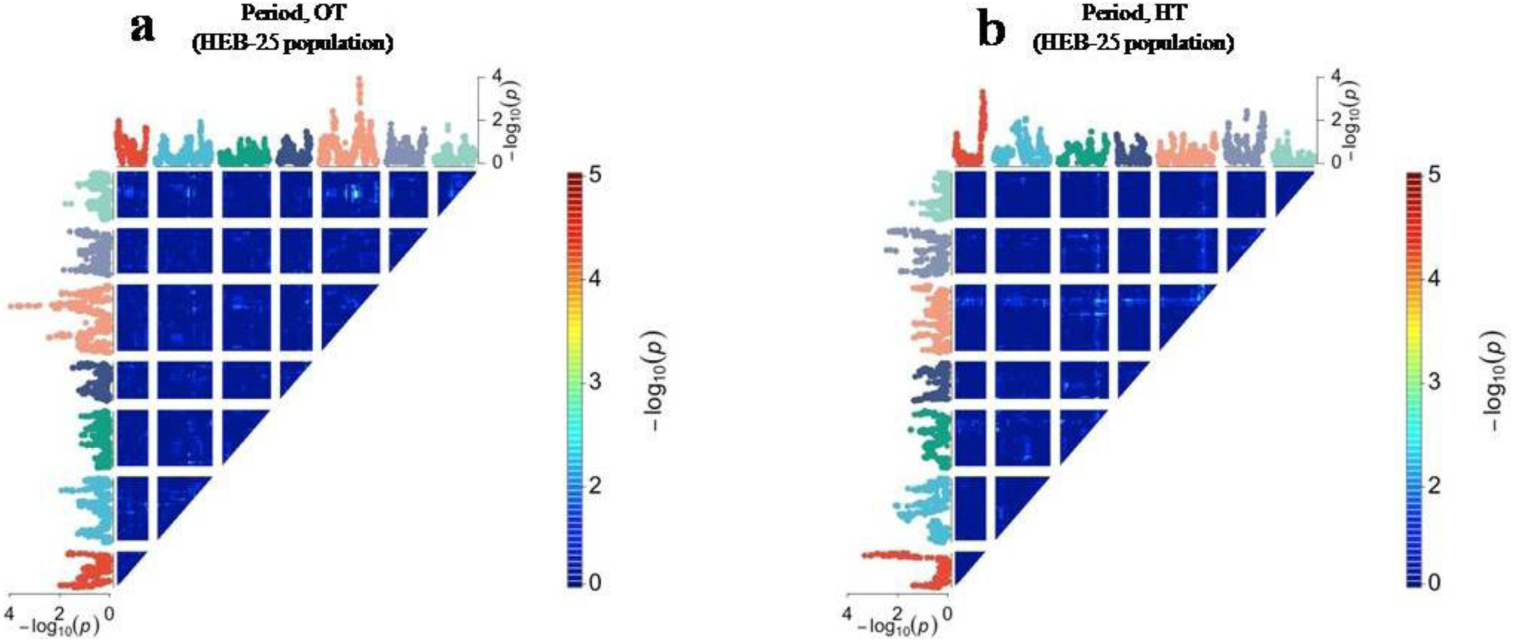
Genome-wide scan for period in HEB-25 population under (a) optimal temperature (OT) and (b) high temperature (HT).Heat map for two-dimensional genome scan with a two-loci interaction model. The Manhattan plots on x- and y-axes in each panel indicate result of one-locus model genome scan. Genome scans were performed using linear mixed model that account population structure as the covariates.

**Figure S6:**
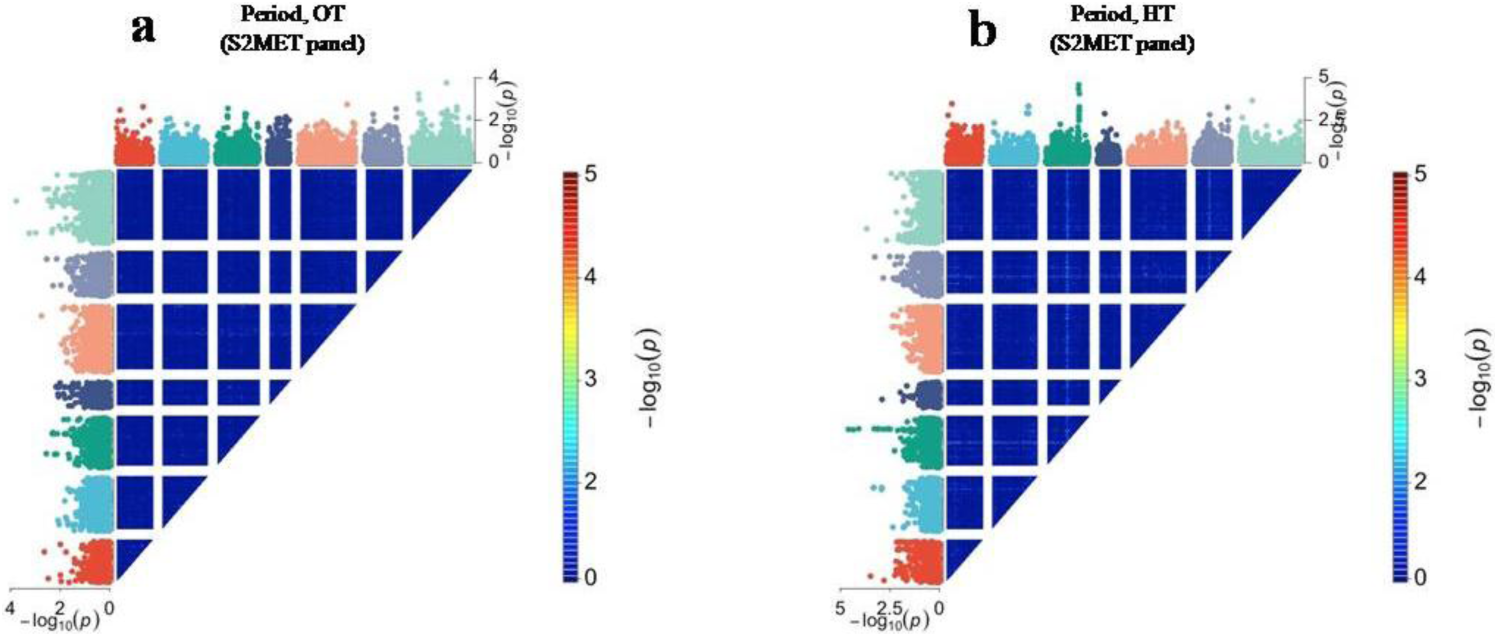
Genome-wide scan for amplitude in S2MET population under (a) optimal temperature (OT) and (b) high temperature (HT).Heat map for two-dimensional genome scan with a two-loci interaction model. The Manhattan plots on x- and y-axes in each panel indicate result of one-locus model genome scan. Genome scans were performed using linear mixed model that account population structure as the covariates.

**Figure S7:**
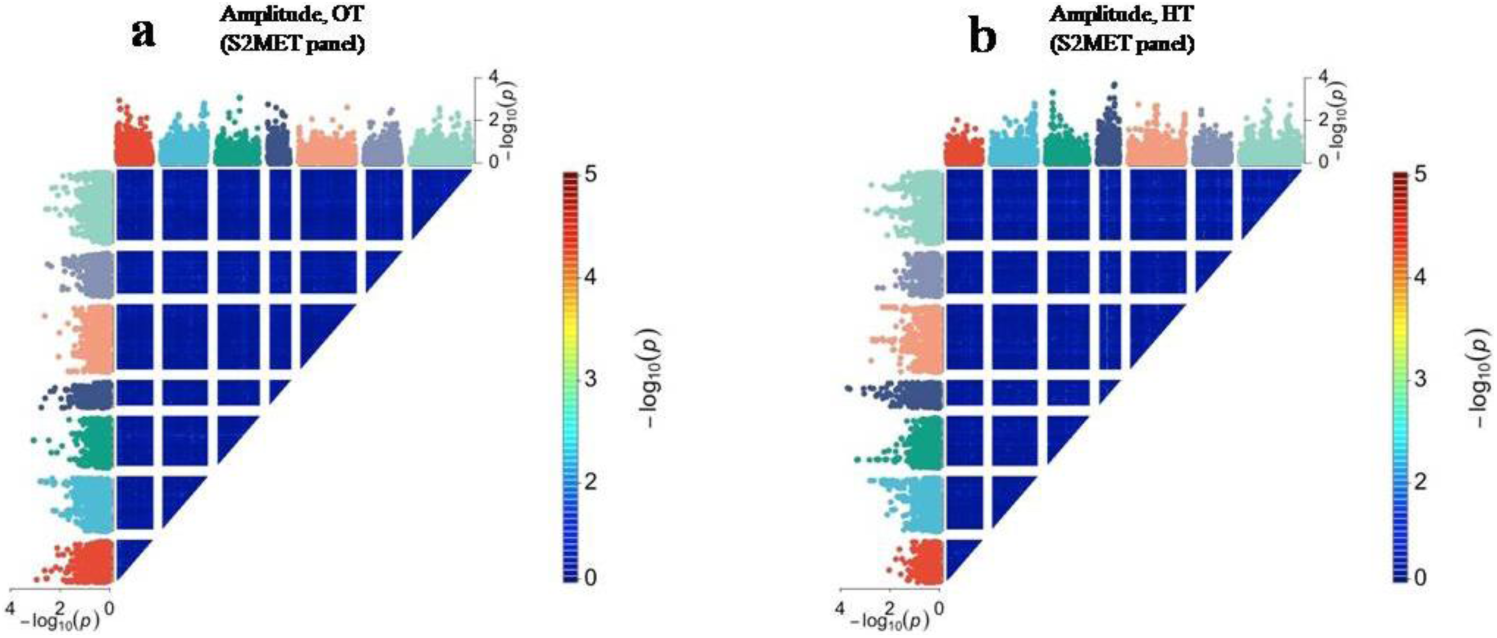
Genome-wide scan for period in S2MET population under (a) optimal temperature (OT) and (b) high temperature (HT).Heat map for two-dimensional genome scan with a two-loci interaction model. The Manhattan plots on x- and y-axes in each panel indicate result of one-locus model genome scan. Genome scans were performed using linear mixed model that account population structure as the covariates.

## Bibliography

1. C. R. McClung, Plant circadian rhythms. Plant Cell (2006) https:/doi.org/10.1105/tpc.106.040980.

2. A. N. Dodd, et al., Cell biology: Plant circadian clocks increase photosynthesis, growth, survival, and competitive advantage. Science (80-.). (2005) https:/doi.org/10.1126/science.1115581.

3. K. Greenham, C. R. McClung, Integrating circadian dynamics with physiological processes in plants. Nat. Rev. Genet. (2015) https:/doi.org/10.1038/nrg3976.

4. K. D. Edwards, J. R. Lynn, P. Gyula, F. Nagy, A. J. Millar, Natural allelic variation in the temperature-compensation mechanisms of the Arabidopsis thaliana circadian clock. Genetics (2005) https:/doi.org/10.1534/genetics.104.035238.

5. W. Y. Kim, et al., ZEITLUPE is a circadian photoreceptor stabilized by GIGANTEA in blue light. Nature (2007) https:/doi.org/10.1038/nature06132.

6. M. K. Choudhary, Y. Nomura, L. Wang, H. Nakagami, D. E. Somers, Quantitative circadian phosphoproteomic analysis of Arabidopsis reveals extensive clock control of key components in physiological, metabolic, and signaling pathways. Mol. Cell. Proteomics (2015) https:/doi.org/10.1074/mcp.M114.047183.

7. Y. Dakhiya, D. Hussien, E. Fridman, M. Kiflawi, R. Green, Correlations between circadian rhythms and growth in challenging environments. Plant Physiol. (2017) https:/doi.org/10.1104/pp.17.00057.

8. E. Bdolach, et al., Thermal plasticity of the circadian clock is under nuclear and cytoplasmic control in wild barley. Plant Cell Environ. (2019) https:/doi.org/10.1111/pce.13606.

9. P. D. Gould, et al., Delayed fluorescence as a universal tool for the measurement of circadian rhythms in higher plants. Plant J. (2009) https:/doi.org/10.1111/j.1365-313X.2009.03819.x.

10. L. Vallelian-Bindschedler, E. Mösinger, J. P. Métraux, P. Schweizer, Structure, expression and localization of a germin-like protein in barley (Hordeum vulgare L.) that is insolubilized in stressed leaves. Plant Mol. Biol. (1998) https:/doi.org/10.1023/A:1005982715972.

11. M. Martínez, I. Rubio-Somoza, P. Carbonero, I. Díaz, A cathepsin B-like cysteine protease gene from Hordeum vulgare (gene CatB) induced by GA in aleurone cells is under circadian control in leaves. J. Exp. Bot. (2003) https:/doi.org/10.1093/jxb/erg099.

12. C. Lillo, Circadian rhythmicity of nitrate reductase activity in barley leaves. Physiol. Plant. (1984) https:/doi.org/10.1111/j.1399-3054.1984.tb05900.x.

13. S. Nagasaka, et al., Time course analysis of gene expression over 24 hours in Fe-deficient barley roots. Plant Mol. Biol. (2009) https:/doi.org/10.1007/s11103-008-9443-0.

14. R. P. Dunford, S. Griffiths, V. Christodoulou, D. A. Laurie, Characterisation of a barley (Hordeum vulgare L.) homologue of the Arabidopsis flowering time regulator GIGANTEA. Theor. Appl. Genet. (2005) https:/doi.org/10.1007/s00122-004-1912-5.

15. A. Turner, J. Beales, S. Faure, R. P. Dunford, D. A. Laurie, Botany: The pseudo-response regulator Ppd-H1 provides adaptation to photoperiod in barley. Science (80-.). (2005) https:/doi.org/10.1126/science.1117619.

16. J. A. Higgins, P. C. Bailey, D. A. Laurie, Comparative genomics of flowering time pathways using brachypodium distachyon as a model for the temperate Grasses. PLoS One (2010) https:/doi.org/10.1371/journal.pone.0010065.

17. C. Campoli, M. Shtaya, S. J. Davis, M. von Korff, Expression conservation within the circadian clock of a monocot: natural variation at barley Ppd-H1 affects circadian expression of flowering time genes, but not clock orthologs. BMC Plant Biol. (2012) https:/doi.org/10.1186/1471-2229-12-97.

18. C. Campoli, et al., HvLUX1 is a candidate gene underlying the early maturity 10 locus in barley: Phylogeny, diversity, and interactions with the circadian clock and photoperiodic pathways. New Phytol. (2013) https:/doi.org/10.1111/nph.12346.

19. S. Faure, et al., Mutation at the circadian clock gene EARLY MATURITY 8 adapts domesticated barley (Hordeum vulgare) to short growing seasons. Proc. Natl. Acad. Sci. U. S. A. (2012) https:/doi.org/10.1073/pnas.1120496109.

20. S. Zakhrabekova, et al., Induced mutations in circadian clock regulator Mat-a facilitated short-season adaptation and range extension in cultivated barley. Proc. Natl. Acad. Sci. U. S. A. (2012) https:/doi.org/10.1073/pnas.1113009109.

21. L. Müller, et al., Differential effects of day-night cues and the circadian clock on the barley transcriptome. Plant Physiol. (2020) https:/doi.org/10.1104/pp.19.01411.

22. C. Bendix, C. M. Marshall, F. G. Harmon, Circadian Clock Genes Universally Control Key Agricultural Traits. Mol. Plant 8, 1135–1152 (2015).

23. R. L. Murphy, et al., Coincident light and clock regulation of pseudoresponse regulator protein 37 (PRR37) controls photoperiodic flowering in sorghum. Proc. Natl. Acad. Sci. U. S. A. (2011) https:/doi.org/10.1073/pnas.1106212108.

24. N. A. Müller, L. Zhang, M. Koornneef, J. M. Jiménez-Gómez, Mutations in EID1 and LNK2 caused light-conditional clock deceleration during tomato domestication. Proc. Natl. Acad. Sci. U. S. A. (2018) https:/doi.org/10.1073/pnas.1801862115.

25. S. HÜbner, et al., Strong correlation of wild barley (Hordeum spontaneum) population structure with temperature and precipitation variation. Mol. Ecol. (2009) https:/doi.org/10.1111/j.1365-294X.2009.04106.x.

26. A. Maurer, et al., Modelling the genetic architecture of flowering time control in barley through nested association mapping. BMC Genomics (2015) https:/doi.org/10.1186/s12864-015-1459-7.

27. J. L. Neyhart, et al., Registration of the S2MET Barley Mapping Population for Multi-Environment Genomewide Selection. J. Plant Regist. (2019) https:/doi.org/10.3198/jpr2018.06.0037crmp.

28. N. A. Müller, et al., Domestication selected for deceleration of the circadian clock in cultivated tomato. Nat. Genet. (2015) https:/doi.org/10.1038/ng.3447.

29. L. Merchuk-Ovnat, et al., Genome scan identifies flowering-independent effects of barley HsDry2.2 locus on yield traits under water deficit. J. Exp. Bot. (2018) https:/doi.org/10.1093/jxb/ery016.

30. E. B. Josephs, Determining the evolutionary forces shaping G × E. New Phytol. (2018) https:/doi.org/10.1111/nph.15103.

31. Y. Shavrukov, et al., Early flowering as a drought escape mechanism in plants: How can it aid wheat production? Front. Plant Sci. (2017) https:/doi.org/10.3389/fpls.2017.01950.

32. K. M. Olsen, J. F. Wendel, A Bountiful Harvest: Genomic Insights into Crop Domestication Phenotypes. Annu. Rev. Plant Biol. (2013) https:/doi.org/10.1146/annurev-arplant-050312-120048.

33. A. L. Caicedo, et al., Genome-wide patterns of nucleotide polymorphism in domesticated rice. PLoS Genet. (2007) https:/doi.org/10.1371/journal.pgen.0030163.

34. S. I. Wright, et al., Evolution: The effects of artificial selection on the maize genome. Science (80-.). (2005) https:/doi.org/10.1126/science.1107891.

35. S. Tanksley, S. McCouch, Seed banks and molecular maps: unlocking genetic potential from the wild. Science (80-.). 277, 1063–1066 (1997).

36. H. Bourguiba, et al., Loss of genetic diversity as a signature of apricot domestication and diffusion into the Mediterranean Basin. BMC Plant Biol. 12, 49 (2012).

37. J. Shi, J. Lai, Patterns of genomic changes with crop domestication and breeding. Curr. Opin. Plant Biol. (2015) https:/doi.org/10.1016/j.pbi.2015.01.008.

38. A. Pankin, J. Altmüller, C. Becker, M. von Korff, Targeted resequencing reveals genomic signatures of barley domestication. New Phytol. (2018) https:/doi.org/10.1111/nph.15077.

39. M. A. Jones, Retrograde signalling as an informant of circadian timing. New Phytol., 1–5 (2018).

40. E. Fridman, M. R. Prusty, A. Faigenboim-Doron, E. Bdolach, Barley Cytoplasmic Multi Parent Population (CMPP) for Studying Evolution of Plasticity Under Domestication in The Allied Genetics Conference, (Genetics Society of America, 2020).

41. S. M. Wadgymar, et al., Identifying targets and agents of selection: innovative methods to evaluate the processes that contribute to local adaptation. Methods Ecol. Evol. (2017) https:/doi.org/10.1111/2041-210X.12777.

42. O. Savolainen, M. Lascoux, J. Merilä, Ecological genomics of local adaptation. Nat. Rev. Genet. (2013) https:/doi.org/10.1038/nrg3522.

43. D. B. Lowry, et al., QTL × environment interactions underlie adaptive divergence in switchgrass across a large latitudinal gradient. Proc. Natl. Acad. Sci. U. S. A. (2019) https:/doi.org/10.1073/pnas.1821543116.

44. S. O. Assanga, et al., Mapping of quantitative trait loci for grain yield and its components in a US popular winter wheat TAM 111 using 90K SNPs. PLoS One (2017) https:/doi.org/10.1371/journal.pone.0189669.

45. A. Börner, G. H. Buck-Sorlin, P. M. Hayes, S. Malyshev, V. Korzun, Molecular mapping of major genes and quantitative trait loci determining flowering time in response to photoperiod in barley. Plant Breed. (2002) https:/doi.org/10.1046/j.1439-0523.2002.00691.x.

46. J. Comadran, et al., Natural variation in a homolog of Antirrhinum CENTRORADIALIS contributed to spring growth habit and environmental adaptation in cultivated barley. Nat. Genet. 44, 1388–92 (2012).

47. T. Zielinski, A. M. Moore, E. Troup, K. J. Halliday, A. J. Millar, Strengths and limitations of period estimation methods for circadian data. PLoS One (2014) https:/doi.org/10.1371/journal.pone.0096462.

48. M. M. Julkowska, S. Saade, G. Agarwal, G. Gao, Y. Pailles, MVApp — Multivariate Analysis Application for Streamlined Data Analysis and Curation 1 [OPEN]. 180, 1261–1276 (2019).

49. C. M. Mutshinda, M. J. Sillanpää, Extended Bayesian LASSO for multiple quantitative trait loci mapping and unobserved phenotype prediction. Genetics (2010) https:/doi.org/10.1534/genetics.110.119586.

50. T. Park, G. Casella, The Bayesian Lasso. J. Am. Stat. Assoc. (2008) https:/doi.org/10.1198/016214508000000337.

51. , VIGoR: Variational Bayesian Inference for Genome-Wide Regression. J. Open Res. Softw. (2016) https:/doi.org/10.5334/jors.80.

52. P. J. Bradbury, et al., TASSEL: Software for association mapping of complex traits in diverse samples. Bioinformatics (2007) https:/doi.org/10.1093/bioinformatics/btm308.

53. C. Xhaard, et al., Heritability of a resting heart rate in a 20-year follow-up family cohort with GWAS data: Insights from the STANISLAS cohort. Eur. J. Prev. Cardiol. (2019) https:/doi.org/10.1177/2047487319890763.

54. H. Perdry, C. Dandine-Roulland, gaston: Genetic Data Handling (QC, GRM, LD, P.A. & Linear Mixed Models.

